# PhyloHerb: A phylogenomic pipeline for processing genome skimming data for plants

**DOI:** 10.1101/2021.11.29.470431

**Authors:** Liming Cai, Hongrui Zhang, Charles C. Davis

**Affiliations:** Harvard University Herbaria, 22 Divinity Avenue, Cambridge, MA 02138, USA; Department of Botany and Plant Sciences, University of California, Riverside, CA 92507, USA; Department of Integrative Biology, University of Texas, Austin, TX 78705, USA

**Keywords:** Herbariomics, high throughput sequencing, mitochondria, plastome, ribosomal genes

## Abstract

- *Premise of the study*: The application of high throughput sequencing, especially to herbarium specimens, is greatly accelerating biodiversity research. Among various techniques, low coverage Illumina sequencing of total genomic DNA (genome skimming) can simultaneously recover the plastid, mitochondrial, and nuclear ribosomal regions across hundreds of species. Here, we introduce PhyloHerb — a bioinformatic pipeline to efficiently and effectively assemble phylogenomic datasets derived from genome skimming.
- *Methods and Results*: PhyloHerb uses either a built-in database or user-specified references to extract orthologous sequences using BLAST search. It outputs FASTA files and offers a suite of utility functions to assist with alignment, data partitioning, concatenation, and phylogeny inference. The program is freely available at https://github.com/lmcai/PhyloHerb/.
- *Conclusions*: Using published data from Clusiaceae, we demonstrated that PhyloHerb can accurately identify genes using highly fragmented assemblies derived from sequencing older herbarium specimens. Our approach is effective at all taxonomic depths and is scalable to thousands of species.

## INTRODUCTION

Herbarium specimens provide the most reliable links between taxonomy, phenotypic traits, genetic information, and species distributions. Beyond their traditional uses, they are increasingly utilized to understand the impacts of global change (Meineke et al., 2018). The advent of high-throughput digitization and industrial scale sequencing of herbarium specimens presents unparalleled opportunities to investigate species diversity in a phylo-spatio-temporal context. Recently, protocols allowing for massive DNA extraction and sequencing of herbarium specimens have been increasingly implemented in large-scale systematic (Nevill et al., 2020; Folk et al., 2021) and ecological investigations (Nitta et al., 2017). These studies often rely on cost-effective library reconstruction and sequencing strategies such as genome skimming, hybrid enrichment, or genotyping by sequencing. Genome skimming (Straub et al., 2012), in particular, is designed to target high-copy conserved regions including plastid, ribosomal, and mitochondrial loci. The streamlined library preparation protocol makes this technique easily automized in wet bench workflows (e.g., robot library preparation). The resulting plastid and ribosomal regions have also provided the bread-and-butter for plant systematics since the 1980s (Palmer and Zamir, 1982), especially for massive scale studies (Zanne et al., 2014; Li et al., 2021). To harvest organelle genomes from short-read data, various assembly software have been developed, including GetOrganelle (Jin et al., 2020), FastPlast (McKain and Wilson, 2017), and NOVOPlasty (Dierckxsens et al., 2017). Annotation tools such as GeSeq (Tillich et al., 2017), Verdant (McKain et al., 2017), and PGA (Qu et al., 2019) have also been implemented in parallel to produce publication quality annotations. However, there are several limitations associated with these annotation tools, hindering their ability to efficiently assemble phylogenetic matrices at a massive scale and for lower-quality data. First, assemblies often return as fragmented scaffolds owing to the degraded herbarium DNA and low base coverage in genome skimming (Forrest et al., 2019). These fragmented assemblies are prone to assembly errors, which may break the synteny of genes, posing additional challenges for accurate annotation. It is thus recommended to align the fragmented assembly against a reference genome to better establish homology (Qu et al., 2019). Second, many of these tools are web-based (e.g., GeSeq and Verdant) or do not allow multiple genomes to be analyzed simultaneously (Table 1). Thus, they cannot support batched analyses over hundreds of species and introduce additional hassles involving data transfer. Third, these existing tools output annotations in GenBank or GFF formats but often require a third-party tool to extract individual genes, further slowing the workflow. Moreover, short tRNA genes and intergenic regions are often excluded from downstream phylogenetic analyses. This is problematic because loci that include such intergenic regions can be more phylogenetically informative (e.g., the *trnL-trnF* spacer).

**Table 1.** Comparison of existing plastome annotation tools. The execution time for PhyloHerb is estimated on the Lenovo SD650 NeXtScale server of the FASRC Cannon compute cluster at Harvard. The execution time for all other software is cited from (Qu et al., 2019).

| Tools | User interface | Time | Output format | Accept multi-FASTA/fragmented assembly |
| --- | --- | --- | --- | --- |
| Plann | Console | ~30s | tbl | No. |
| Verdant/annoBTD | Web/Console | 10–30 min | GFF3 | Currently not supported but can be incorporated (pers. comm. with author). |
| GeSeq | Web | 6 s–13 min | GenBank | Yes. One assembly per run for fragmented genomes. |
| PGA | Console | ~20s | GenBank | Yes, but not recommended. Batch processing. |
| PhyloHerb | Console | 2–30s | FASTA | Yes. Batch processing. |

To bridge this impasse, we present PhyloHerb, a command line tool for the simultaneous annotation, alignment, and phylogenetic estimation of thousands of species using genome skimming data. The core function of PhyloHerb is to apply BLAST searches to identify locus boundaries using the built-in database (GeneBank reference annotations) or customized references. This allows a user to extract gene and intergenic regions *en masse* from assemblies with marginal quality and directly output orthologous sequences into FASTA format for downstream phylogenetic analyses. It also offers a host of functionality to assist with genome assembly, evaluate assembly statistics, concatenate loci, generate gene partition file, and curate alignment for easier manual inspection. Our lab has been applying this tool to assemble phylogenetic datasets for published and ongoing systematic studies in flowering plants and algae (Marinho et al., 2019; Lyra et al., 2021). These published and ongoing datasets include more than 1,500 species with a median base coverage 21.9x for the plastid genome. Within less than one hour of CPU time, PhyloHerb can compile orthologous FASTA sequences for one thousand species across 150 genetic blocks in the plastid, nuclear, and mitochondrial genome. We also piloted this tool recently with a group of scientists, including many first-time users, at a day-long workshop hosted by our authorship team at the 2021 Botany meeting (https://2021.botanyconference.org/engine/search/index.php?func=detail&aid=20).

## METHODS AND RESULTS

PhyloHerb is an open-source program (GNU General Public License) written in Python 3. The source code, user manual, as well as the example dataset, are freely available at https://github.com/lmcai/PhyloHerb/. The software can be easily installed on linux, OSX, and Windows systems by simply decompressing the source code package. Before implementing the software, users need to install the python modules Biopython and ete3, and BLAST+ (Johnson et al., 2008). Specific installation instructions can be found on the Github website (https://github.com/lmcai/PhyloHerb/tree/main/botany2021_tutorial).

### Input preparation

The minimum input for PhyloHerb includes the raw assemblies of plastid, ribosomal, and mitochondrial genomes in FASTA format (Fig. 1). We recommend GetOrganelle (Jin et al., 2020) for *de novo* assembly of these three genomes, which has been demonstrated to be state-of-the-art for this initial step (Freudenthal et al., 2020). Once assemblies are obtained, users can use the ‘qc’ function in PhyloHerb to evaluate the quality of their assemblies. When providing assemblies alone, this function will generate a summary spreadsheet for the following information: assembly size in base pairs (bp), number of scaffolds, and GC content. If the assembly is generated from GetOrganelle, PhyloHerb will read the log files from GetOrganelle and output the following additional statistics: total input reads, number of reads in the target region, average base coverage, and whether the genome is circularized.

**Figure 1.**
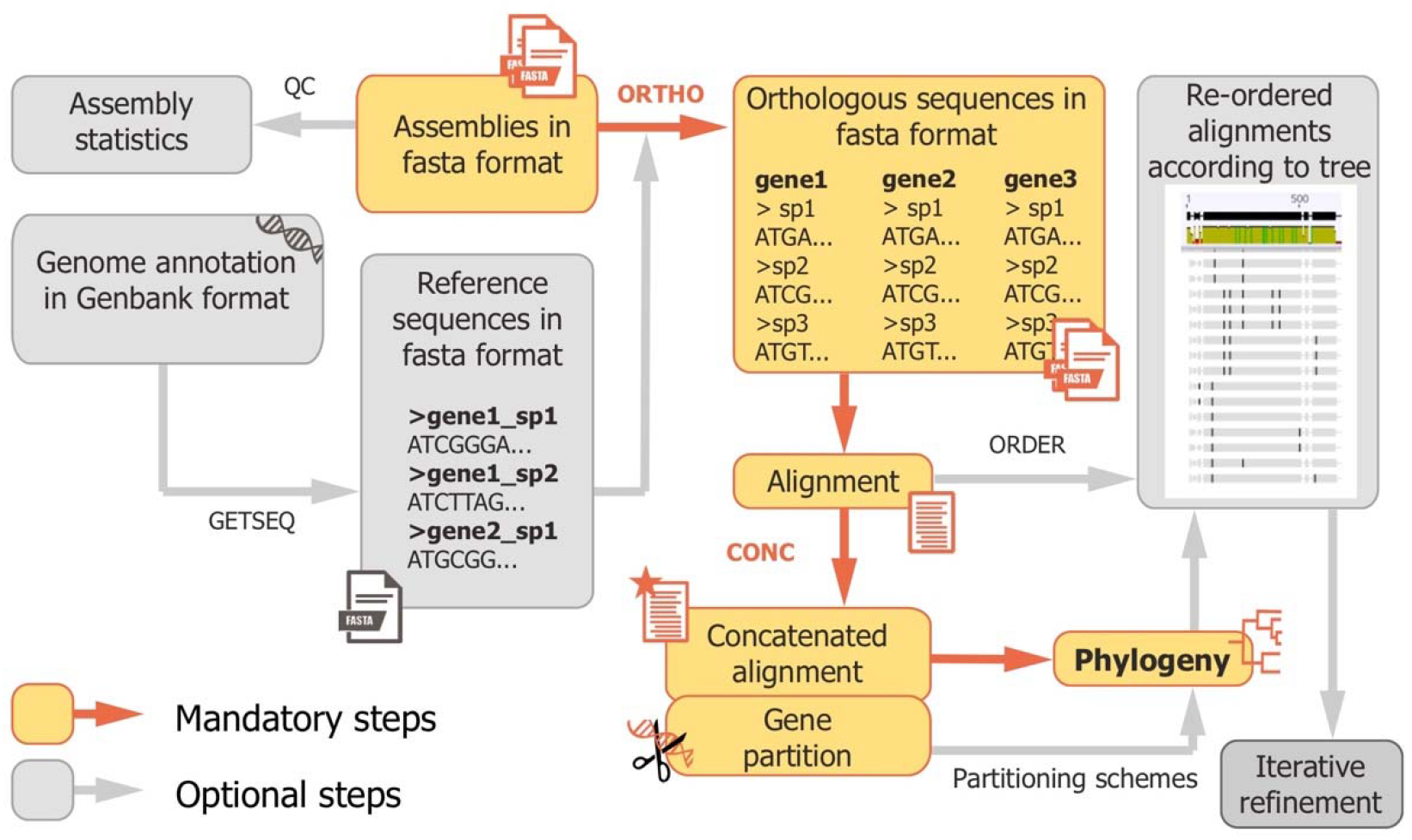
PhyloHerb workflow. The five main function modules of PhyloHerb, including qc, getseq, ortho, conc, and order, provide a versatile and efficient tool to curate and analyze genome skimming data.

PhyloHerb relies on a reference database to identify orthologs using BLAST searches. A built-in reference plant database is included in the source code and is explained in detail below. This database is comprehensive for land plant organellar genes and the nuclear rRNA repeat region. Users can also specify their own references to define loci. This user specified reference should be a single FASTA file containing sequences from all targeted loci. The sequence header should start with the locus name, followed by an underscore ‘_’, and any additional characters to distinguish different copies (Fig. 1). In the following conditions users may consider using their own references: 1) Consolidate multiple short gene and intergenic regions to a longer locus for better BLAST results; 2) Use closely related species for more accurate annotation; 3) Harvest loci not included in the database. For example, PhyloHerb can be used to extract *LFY* — a single-copy nuclear phylogenetic marker (Frohlich and Meyerowitz, 1997) — from transcriptome assemblies when provided with a *LFY* reference sequence.

### Locating locus boundaries

The ‘ortho’ function of PhyloHerb uses a reverse query-subject BLAST approach to locate loci within an assembly (Qu et al., 2019). Here, the BLAST database is constructed from the unannotated assemblies, while the reference nucleotide database is used for BLAST queries. The input files are genome assemblies in FASTA format, and the outputs are FASTA files of individual loci. Our built-in plastid database referenced above contains 98 genes (Appendix S1) from 355 land plants (Appendix S2). To prepare this plastid database, we downloaded all available plastid genome annotations in GenBank (accessed June 17, 2021) and selected one representative species per family. Therefore, the plastid genes in the database represents the union of all identified protein coding genes and rRNAs in land plants (bryophytes, ferns, gymnosperms, and angiosperms). We also manually curated the database to correct synonymous gene and annotation errors. The mitochondrial and nuclear rRNA database were prepared similarly. The mitochondrial database includes 71 genes (Appendix S3) from 68 species (Appendix S4). The nuclear rRNA database of 18S, 28S, and 5.8S includes 155 species (Appendix S5). For each locus, reference sequences from all species in the database will be blasted to the unannotated assembly with an e-value threshold of 1e-20 and length threshold of 60 bp. The BLASTN hit with lowest e-value and longest alignment length will be used to establish the boundaries of genes. A subset of genes and species can be included in the analysis by invoking the ‘-sp’ and ‘-g’ flags, respectively. The minimum length threshold can be adjusted using the ‘-l’ flag and the minimum e-value can be modified using the ‘-evalue’ flag. For the two internal transcribed spacers (ITSs), the gene locations of the three rRNAs are used to identify the start and end site of ITS. Here, we use gene synteny instead of sequence similarity to avoid spurious BLAST hits associated with high sequence divergence. The external transcribed spacer (ETS) and non-transcribed spacer (NTS) will not be automatically extracted by PhyloHerb.

To obtain intergenic regions, PhyloHerb uses BLASTN search instead of gene synteny despite high sequence variation. This is because structural changes and fragmented assemblies can confound ortholog identification in our empirical practices. To more accurately determine locus boundaries, we recommend using closely related species as references and including conserved gene regions on both ends to define locus boundaries. For example, twelve short genes, including *psbJ, psbL*, and *rpl20*, are arranged linearly in a 5 kb block in the plastid genome of *Arabidopsis thaliana* (Fig. 2A). To include intergenic regions, we can group these loci into two segments, each approximately 2.5 kb in length and containing five to seven genes (LOC 1 and 2 in Fig. 2A). Here, the defined genetic block should not exceed 5 kb in length for closely related species (e.g., species in closely related genera) and 3 kb for more divergent lineages. This is because structural changes can break the synteny of the genome, leading to truncated BLAST hits. Once the boundaries of genetic blocks are defined, PhyloHerb offers the function ‘getseq’ to extract corresponding regions from GenBank formatted genome annotations. The input files include a genetic block definition file and GenBank annotations (Fig. 2B). This function outputs FASTA files of the designated regions that can be directly used as reference in the ‘ortho’ function. The ‘getseq’ function offers two modes ‘genetic_block’ and ‘intergenic’, which will include gene sequences on both ends or include intergenic regions only, respectively (Fig. 2B). The ‘genetic_block’ mode is suitable for getting longer loci spanning multiple genes. We anticipate that this later functionality will be especially relevant for resolving clades at shallower phylogenetic depths. The ‘intergenic’ mode can be used to obtain the intergenic regions between two adjacent genes.

**Figure 2.**
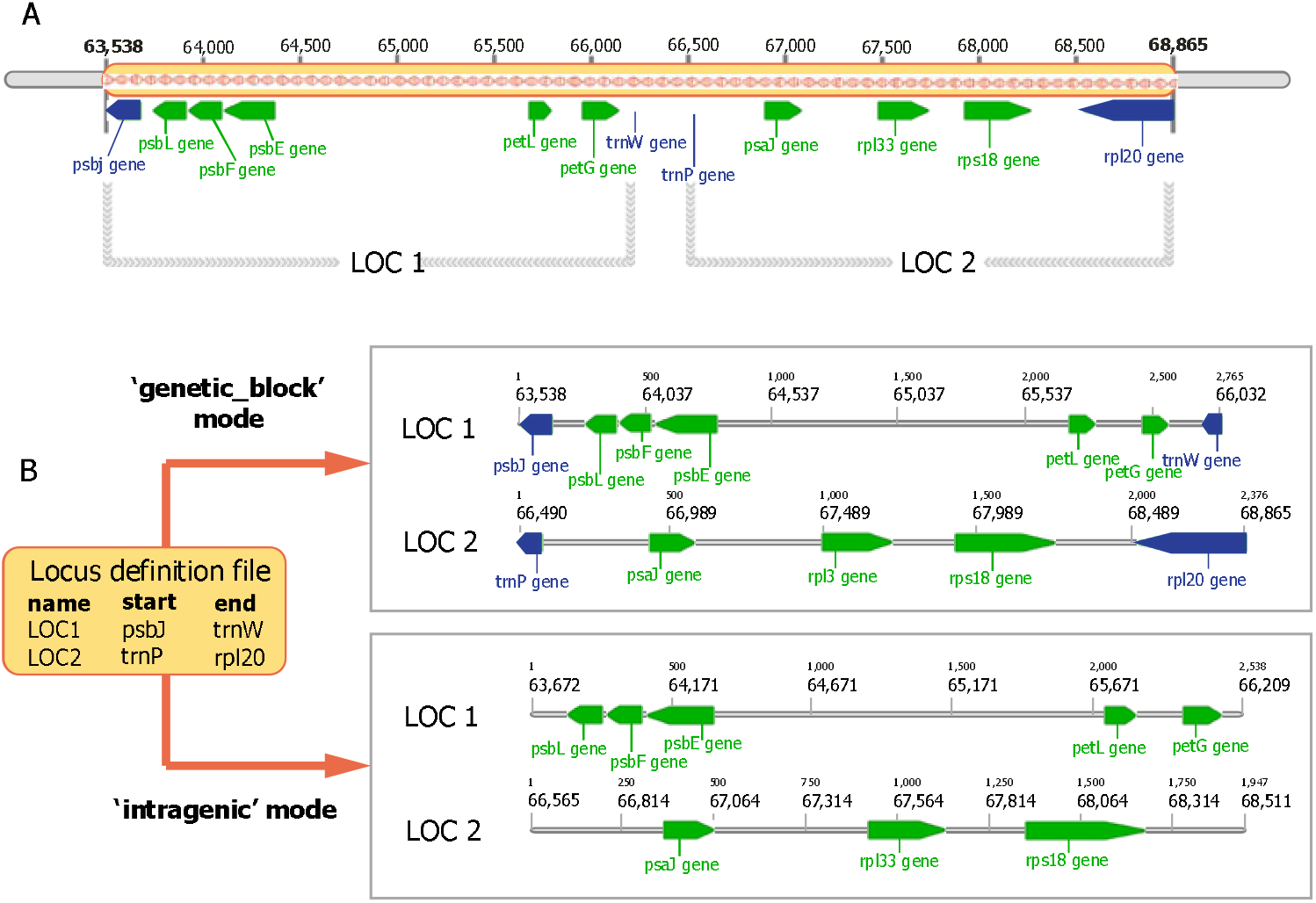
Defining and extracting genetic block with PhyloHerb. (A) A 5 kb long continuous genetic block on the plastid genome of *Arabidopsis thaliana* divided into two loci (LOC1 and LOC2). (B) The ‘getseq’ function of PhyloHerb can be used to extract sequences of predefined genetic blocks. The ‘genetic_block’ mode will include genes on both ends, while the ‘intergenic’ mode does not.

### Utility functions for phylogenetic analysis

PhyloHerb offers several useful utility functions to assist alignment and phylogenetic reconstruction. We recommend MAFFT (Katoh and Standley, 2013) for aligning conserved genes, and PASTA (Mirarab et al., 2015) for aligning more variable intergenic regions. An example bash file is provided in our source package (https://github.com/lmcai/PhyloHerb/blob/main/phyloherbLib/mafft_pasta.sh). Once individual alignments are generated, users can implement the ‘conc’ function in PhyloHerb to concatenate sequences. The list and order of loci to be included can be customized using the ‘-g’ flag. The ‘conc’ function is especially suitable for creating large matrices with hundreds of species and genes, for which other GUI applications, such as MEGA (Tamura et al., 2007), often suffer from insufficient memory. PhyloHerb will also generate a gene partition file that can be directly input into PartitionFinder (Lanfear et al., 2017). The inferred partition scheme and the concatenated sequences can then be applied to phylogeny inference tools such as RAxML (Stamatakis, 2014), IQ-TREE (Minh et al., 2020), or ExaML (Kozlov et al., 2015).

Large-scale phylogenetic studies often require iterative alignment–phylogeny refinement practices to clean sequence data. To do so, researchers often visualize and edit the alignments in tools such as Geneious (https://www.geneious.com). Here, reordering sequences according to the species tree can help distinguish shared mutations between close relatives versus spuriously aligned regions arising from assembly or BLAST errors. Therefore, we developed the ‘order’ function of PhyloHerb, which takes a reference tree and reorders all input alignments based on the phylogeny (Fig. 1). It also offers the option to remove sequences with excessive missing data via the ‘-missing’ flag. A float number from 0 to 1 can be used to indicate the maximum proportion of ambiguous sites allowed for each sequence. This function will generate an ordered alignment and a pruned species tree for each locus. The pruned species tree can be used to guide the PASTA alignment in the second round, which will significantly improve the alignment of especially intergenic regions (Mirarab et al., 2015).

### Example workflow applied to Clusiaceae

We selected ten species from three genera in the Clusiaceae family from the published dataset by Marinho et al. (Marinho et al., 2019; Table 2) to verify the utility of our pipeline. The input data was generated from paired end Illumina Hi-Seq 2×125 sequencing platform and their size ranged from 81.5 Mb to 517.8 Mb. All testing was conducted on the Lenovo SD650 NeXtScale server of the FASRC Cannon compute cluster at Harvard. A detailed tutorial is provided in Appendix S6.

After generating genomes using GetOrganelle (Jin et al., 2020), we used the ‘qc’ function of PhyloHerb to summarize the assembly statistics, which took 0.53s CPU time for ten species. The plastid genome assemblies ranged from 128.2 kb to 164.3 kb (Appendix S7). Three of ten species have fully circularized plastid genomes. The coverage of rRNA is generally higher than the plastid organelle and seven species have complete rRNA repeats assembled. The mitochondria assemblies are more fragmented, ranging from 6.5 to 359.3 kb in size. We then used the ‘ortho’ function of PhyloHerb to extract orthologous regions. For ten species, this took 297.7s CPU time and 159 Mb peak memory using the built-in plastid database. When using a custom reference from *Garcinia gummi-gutta* (Clusiaceae, GenBank accession number NC_047250.1), only 6.6s CPU time and 153 Mb peak memory was needed. For nuclear rRNA, this step took 2.6s CPU time and 47 Mb peak memory. For mitochondrial genes, it took 17.4s CPU time and 108 Mb peak memory. In the resulting sequence matrices, no missing data are found in the plastid and nuclear ribosomal genes. But the 53 mitochondrial genes contained from 0 to 90% missing data, reflecting the lower assembly quality of mitochondrial genomes. Compared to the GenBank reference annotation of *G. gummi-gutta*, PhyloHerb recovered all 81 plastid genes in the reference, and six additional genes. Five of these six genes (*psbG, ycf10, ycf15, ycf68*, and *ycf9*) were not annotated in the reference but do exist in the plastid genome, but the *rpl32* gene was indeed mis-specified by PhyloHerb. Here, all species had a false positive *rpl32* BLAST hit in their *rpl23* region, while *rpl32* was absent in Clusiaceae. Using gene tree phylogeny, we confirmed that no paralogous copies were included in any of the 87 plastid genes, including the *rpl32* gene (Appendix S8). Therefore, the incorrect gene assignment does not introduce bias in phylogenetic inference in this case. We anticipate that shorter genes will have higher risks of such mis-assignment, but this is not unique to our pipeline (Qu et al., 2019). To reduce this possibility, we recommend consolidating multiple adjacent short genes into longer genetic blocks to mitigate false positive BLAST hits and to include informative intergenic regions (see Section ‘Intergenic region improves phylogenetic resolution’ below).

To infer a species tree based on the plastid genes, we used MAFFT v7.407 (Katoh and Standley, 2013) for alignment and then used the ‘conc’ function in PhyloHerb to concatenate individual alignments. This concatenation step took 5.8s CPU time with 64 Mb peak memory for 87 plastid genes across ten species. Finally, we used IQ-TREE v2.0.5 (Minh et al., 2020) to infer a species tree based on the concatenated alignment using a GTRGAMMA substitution model and 1000 ultrafast bootstrap replicates (UFBP). All internal nodes were maximumly supported except for *Tovomita acutiflora* and *Tovomita choisyana*, which only received 45 UFBP (Fig. 3A).

**Figure 3.**
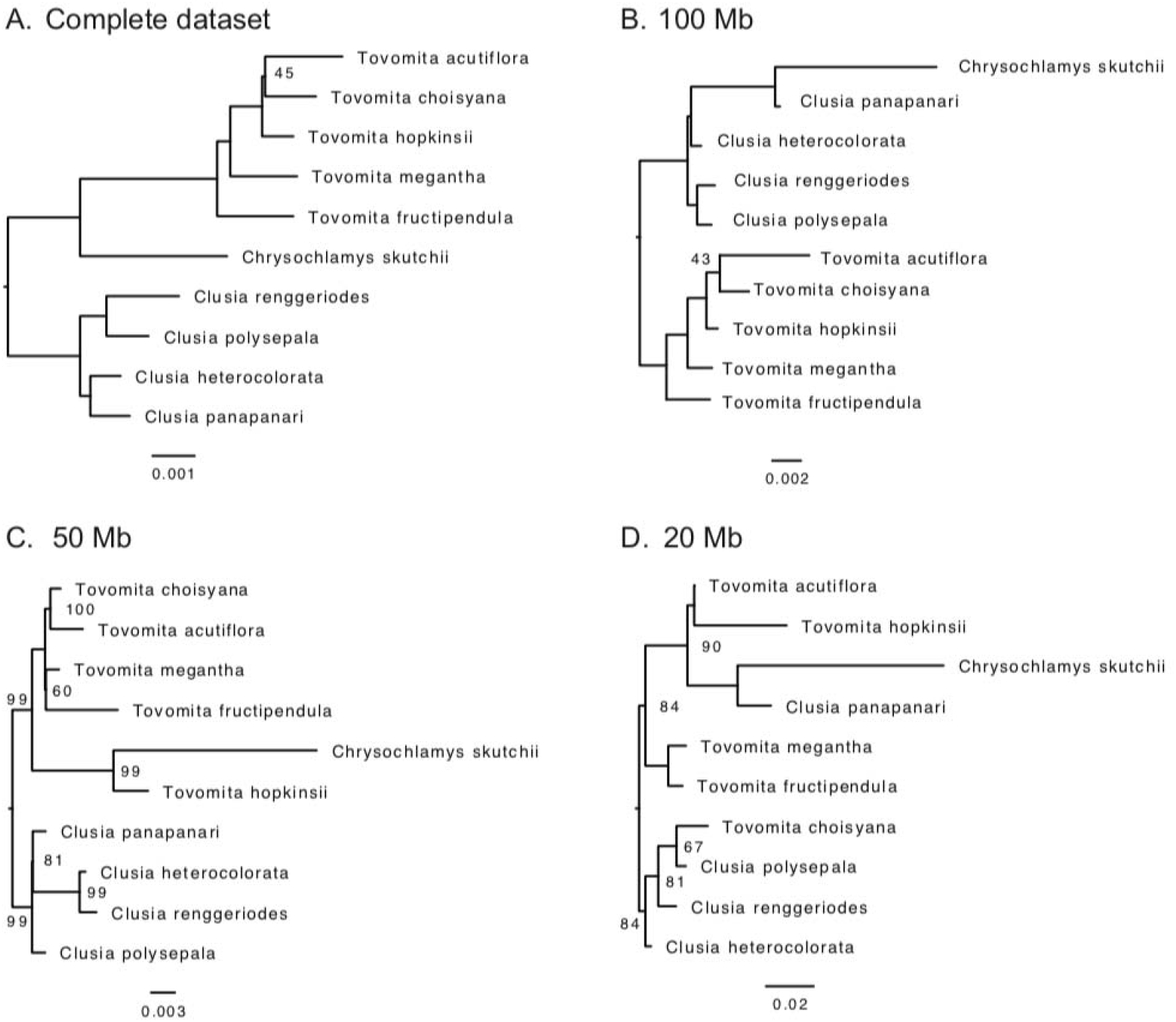
Phylogeny of ten Clusiaceae species inferred from the complete and subsampled plastid datasets. The size of the input raw read data is presented on the upper left of each panel. Nodal support was estimated from 1000 ultrafast bootstrap (UFBP). Only numbers smaller than 100 UFBP were displayed. For all four analyses, an unpartitioned concatenated DNA alignment of 87 plastid genes was used to infer the species tree in IQ-TREE using the GTR+GAMMA model.

### Scalability

The two most resource-consuming steps in PhyloHerb are ortholog extraction and alignment concatenation. To demonstrate its scalability to massive datasets, we applied it to our unpublished dataset from an ongoing project focusing on Malpighiales. We randomly selected 1,000 species from this larger dataset. These species share a common ancestor at about 90 million years ago (Xi et al., 2012) and the average pairwise sequence divergence is 0.035 for plastid genes. We used a single outgroup species *Vitis vinifera* (GenBank accession number: NC_007957.1) as reference to extract plastid genes. It took 357.1s CPU time (~2.5 h wall clock time for a single thread) with a peak memory of 53 Mb to extract all 84 plastid genes in 1000 assemblies. After aligning each of the gene sequences with MAFFT, we used the ‘conc’ function of PhyloHerb to create a concatenated matrix of 247 kb aligned sites across 1,000 species. This step took 1127.4s CPU time with 4.43 Gb peak memory. We have attempted to apply this dataset to other popular programs that offers concatenation functions including Geneious, MEGA (Tamura et al., 2007), and SeaView (Gouy et al., 2010). Both MEGA and SeaView crashed shortly after we initiated the concatenation on our laptop (MacBook Pro 2.5 GHz Intel Core i7 with 16 Gb RAM). In contrast, Geneious was capable of concatenated this large matrix, which took 158.23s CPU time and 2.14 Gb peak memory on the MacBook. But according to the user manual (https://www.geneious.com/), Geneious is not designed to work efficiently with more than 10,000 sequences (e.g., 100 loci x 100 species). Moreover, this concatenation function is also proprietary in Geneious and requires a paid subscription, which creates barriers for accessibility.

### Intergenic regions improve phylogenetic resolution

For rapidly diverging lineages, coding regions may contain little information to fully resolve the phylogeny (Gielly and Taberlet, 1994). In these cases, intergenic regions bring additional informative sites to improve phylogenetic resolution. To demonstrated this, we compared the phylogenetic trees of ten Clusiaceae species built from gene regions alone versus gene and intergenic regions. We used the ‘getseq’ and ‘ortho’ function of PhyloHerb to obtain sequences of four plastid genetic blocks. Each genetic block contained five to seven short genes and ranged between 2.3 to 3.7 kb (Appendix S9). The coding regions accounted for 37.0% of the total length, but only 10.3% of the phylogenetic informative sites. Therefore, most phylogenetic informative sites reside within the intergenic regions. Consequently, the species tree inferred from both gene and intergenic regions had higher average nodal support of 93 UFBP compared to the species tree inferred from gene regions only (69 UFBP, Appendix S10). This result demonstrated that the integration of intergenic regions effectively improves phylogenetic resolution by adding more rapidly evolving sites. One caveat here is that establishing site homology among highly variable intergenic regions is especially challenging when sampling deep and shallow phylogenetic depths simultaneously. In such cases, a profile alignment approach, which builds alignment subsets among closely related taxa and then merges them into a single aligned matrix, can be implemented (e.g., PASTA; Mirarab et al., 2015).

### Risks of low coverage

To explore the limits of genome skimming techniques and artifacts attributed to low sequence coverage, we subsampled the Clusiaceae genome skimming data to include only 100 Mb, 50 Mb, and 20 Mb reads. These data translate to an average base coverage of 8.4x, 4.5x, and 2.2x for the plastid genome, respectively (Appendix S11). We applied the same genome assembly and gene extraction pipeline described above. The concatenated matrix from these three datasets were similar in length (approximately 96 kb), but contained 14%, 29%, and 56% ambiguous characters, respectively. The quality of the alignments varied significantly across these subsampled datasets. When we manually inspected in Geneious, the alignment from the complete dataset required minimum adjustment with high sequence identity throughout (Appendix S12). When the input data size was reduced to 100 Mb, we noticed more incidences of assembly or annotation errors requiring removal (green bars in Appendix S12). These biases, combined with the increasing amount of missing data, also reduced the sequence identity significantly. The same trend also applied to the 50 Mb and 20 Mb datasets. Without correction, species phylogenies built from these alignments have incorrect topologies and potentially spurious branch length distributions (Fig. 3). In particular, species with excessive missing data often exhibit long branches (e.g., *Chrysochlamys skutchii* in Fig. 3 and Appendix S11). Based on these results, we empirically conclude that 50 Mb or 5x coverage is the lower limit for plastid genome assembly using genome skimming technique. For such datasets, researchers need to apply more stringent filtering criteria, such as smaller BLAST e-value threshold and using closely related custom references for accurate ortholog assignment.

## CONCLUSIONS

As herbarium specimen sequencing and plastid genome based phylogenomics become increasingly popular and greatly scalable in biodiversity research, bioinformatic tools need to be developed in parallel to accommodate datasets of various size and quality. To identify and extract orthologs for phylogenomic analyses, researchers typically need to annotate the assemblies using standalone software and then extract target loci from the resulting GenBank or GFF annotations. Working with fragmented assemblies like those yielded from degraded herbarium materials is greatly challenging owing to currently unsupported multi-FASTA file formats. PhyloHerb offers an easy-to-use tool that allows users to more effectively and efficiently analyze assemblies of marginal quality. It directly outputs orthologous sequences in FASTA format that can be used for downstream alignment and phylogenomic inference. Users can create custom references to extract intergenic regions, which will likely be crucial to resolve rapidly diverging lineages. However, datasets with fewer than 50 Mb input reads should be processed very carefully because we demonstrated via data subsampling simulations that degraded datasets contain excessive assembly errors and require substantial manual cleaning. In addition, it should be noted that PhyloHerb is not designed to generate polished genome annotations but rather to generate alignments for phylogenetic purposes, which complement the functions of existing tools such as GeSeq or PGA. In PhyloHerb, the accuracy of locus boundary determination is tied with the performance of BLAST searches. For conserved gene regions, PhyloHerb can confidently identify their locations given our comprehensive built-in database across land plants. But for lineages or loci with high sequence divergence, we strongly recommend applying more closely related taxa as references. Moreover, genome structural modifications such as insertion, deletion, and reversion will greatly impact the performance of PhyloHerb, especially when extracting genetic blocks spanning several genes. Where possible, we recommend checking gene synteny using completely circularized plastid genomes to avoid using regions prone to macrostructural change. Even in cases where gene synteny is conserved, the custom genetic loci should not exceed 5 kb or span more than ten genes to avoid truncated BLAST hits. Compared to other annotation tools, PhyloHerb offers additional flexibly in input data quality while demonstrate high annotation accuracy. It represents a powerful and freely available tool that allows researchers to rapidly assemble orthologous alignments from three cellular genomic components.

## ACKNOWLEDGMENTS

We especially thank the participants of workshop we coordinated at Botany 2021 meeting (https://2021.botanyconference.org/engine/search/index.php?func=detail&aid=20) who helped to prototype this tool and provided useful feedback to improve the software. We want to thank Dr. Lucas C. Marinho for revising the figures. We want to thank the members of the Davis lab for insightful comments of the software and manuscript. Startup funds from Harvard University to C.C.D. helped to facilitate this study.

## DATA ACCESSIBILITY

The PhyloHerb source code, as well as the example dataset used to demonstrate its utility, are available at https://github.com/lmcai/PhyloHerb.

## APPENDIX

**Appendix S1** List of 98 plastid genes included in the PhyloHerb built-in reference database.

**Appendix S2** Taxonomic information and GenBank accession number of 355 land plants included in the PhyloHerb built-in plastid reference database.

**Appendix S3** List of 71 mitochondrial genes included in the PhyloHerb built-in reference database.

**Appendix S4** Taxonomic information and GenBank accession number of 68 land plants included in the PhyloHerb built-in mitochondrial reference database.

**Appendix S5** List of 155 land plants included in the PhyloHerb built-in ribosomal gene reference database.

**Appendix S6** Example PhyloHerb workflow applied to Clusiaceae

**Appendix S7** Summary statistics of the organellar and ribosomal assemblies from ten Clusiaceae species

**Appendix S8** Input and output files of PhyloHerb applied to the Clusiaceae datasets.

**Appendix S9** Gene content of four pre-defined loci L1-L4

**Appendix S10** Intergenic regions in the plastid genome improves phylogenetic resolution

**Appendix S11** Plastid genome assembly statistics in the subsampled datasets

**Appendix S12** Increasing risks of systematic errors stemming from low coverage data

